# A low-adhesion mechanism enables axon growth in the developing brain

**DOI:** 10.64898/2026.10.05.756689

**Authors:** Eissa Alfadil, Severin Filser, Benjamin Escribano, Matthias J. Hackl, Frank Bradke

## Abstract

During development, axons grow through complex tissue environments to wire the brain^1^. Axon growth speeds measured in cell culture are incompatible with the distances axons must cover *in vivo*^2,3^. Yet how axons achieve the growth speeds required to establish long-range connectivity within a narrow embryonic developmental window, and by what mechanism, remains unknown. Here, using long-term live imaging in brain sections, mouse genetics, molecular and pharmacological perturbations, together with AxoVivo, a newly developed intravital imaging platform that resolves subcellular structures, including growth cones and cytoskeletal dynamics, in living mouse embryos, we show that axons of cortical projection neurons (CPNs) employ a low-adhesion mechanism to grow. Even as growth cones change shape and turn, axons grow an order of magnitude faster than *in vitro*. The actin motor myosin II controls the growth cone actin cytoskeleton and its morphology but not axon growth itself, which is instead promoted by microtubule stabilisation. Finally, super-resolution shadow imaging (SUSHI)^4^ revealed that growth cones do not pull on the surrounding extracellular matrix. Taken together, this low-adhesion mechanism driven by protruding microtubules ensures the rapid growth necessary for long-distance wiring of the developing nervous system.

## Introduction

During embryonic development, the axons of layer 2/3 cortical projection neurons (CPNs) traverse several millimetres of tissue within days to establish callosal connectivity^2,5,6^. *In vitro*, cortical and hippocampal neurons extend their axons at only 100–150 µm per day^3,7,8^, more than an order of magnitude slower than would be required *in vivo*. This raises the possibility that mechanisms found in culture may inadequately describe the physiologically relevant mode of axon growth. This concern is not unique to neurons: non-neuronal cells, including leukocytes, have been shown to employ fundamentally different modes of motility in the native tissue environment compared to *in vitro*, most notably the capacity for rapid, adhesion-independent migration *in vivo* that is not observed on flat substrates^9^.

In two-dimensional (2D) culture models, it was proposed that axons grow using mechanisms analogous to migrating fibroblasts^10^. This mesenchymal mode of migration is characterised by integrin-dependent adhesion to the extracellular matrix (ECM) and actomyosin-generated pulling forces to enable forward movement^11–13^. In this model, the growth cone acts as a ‘fibroblast on a leash’, pulling itself forward along the ECM and tugging the axon behind. This view was supported by *in vitro* experiments showing that pulling the axon at the growth cone with a glass needle leads to axon extension^14^. However, in three-dimensional (3D) cultures, cortical and hippocampal neurons grow their axons without pulling on the collagen matrix^7^. How axons grow in their natural habitat, and at what rate, has remained unclear.

Here, we investigated the mechanisms of axon growth of CPNs in the developing brain, both *in situ* in brain slices and in living mouse embryos. We found that the physiological growth mode enables axons to grow an order of magnitude faster than in culture, irrespective of the region of the trajectory. Combining live imaging, super-resolution shadow imaging (SUSHI), and pharmacological and molecular perturbations revealed that axon growth is driven by microtubule stabilisation independently of actomyosin-mediated pulling on the ECM. Demonstrating this low-adhesion mode of growth and characterising axon dynamics resolves how the remarkable axon growth speeds *in vivo* implement brain wiring during embryonic development.

## Results

### Axons *in situ* grow rapidly across cortical regions

To investigate the intrinsic mechanisms of axon growth in their native environment, we focused on layer 2/3 CPNs for three reasons: first, CPNs extend axons over long distances and across multiple tissue territories to establish cortico-cortical connections^5^ (**Fig. 1a**). Second, key cytoskeletal and mechanical aspects of axon growth have been extensively characterised in cortical neurons, as well as in hippocampal neurons, in 2D and 3D culture systems^3,7,15–19^, providing a well-defined *in vitro* baseline against which *in situ* and *in vivo* mechanisms can be compared. Third, their entire axon trajectory can be followed *in situ* using a well-established slice culture model^20,21^, which we optimised for high-resolution and long-term imaging of axon development^22^.

**Fig. 1:**
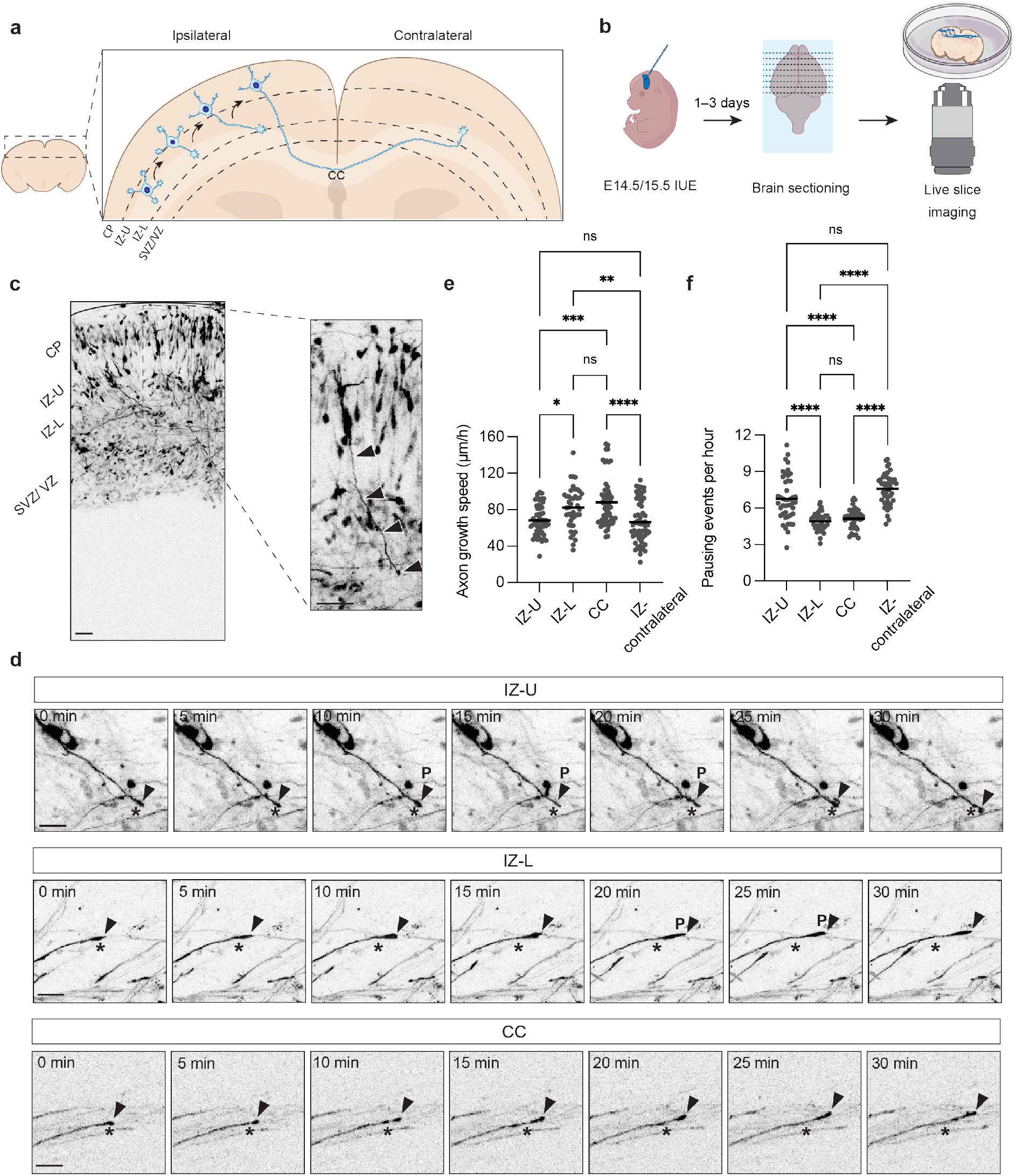
Axon growth speed and pausing are regulated in a region-specific manner. a. A schematic of neuronal polarisation in the mouse cortex. CP, cortical plate; IZ-U, upper intermediate zone; IZ-L, lower intermediate zone; SVZ/VZ, subventricular zone/ventricular zone; CC, corpus callosum. b. A schematic of the pipeline used to visualise axons *in situ,* which included bilateral *in utero* electroporation (IUE), brain sectioning, and live slice imaging. c. A representative tiled image of cortical projection neurons (CPNs) *in situ* (left) and a magnification of an extending axon (right). Neurons are labelled with Lyn-mNeonGreen via IUE. Arrowheads mark the axon. Scale bars, 50 µm. d. Representative images from tiled time-lapses showing axon extension and pausing “P” in the IZ-U, IZ-L and CC. Asterisks mark the initial position of the axon tip. Arrowheads track the moving axon tip. Neurons are labelled with Lyn-mNeonGreen via IUE. Scale bars, 20 µm. e. Quantification of axon speed at different anatomical regions. N = 3 independent experiments, n = 52 axons (IZ-U), n = 43 axons (IZ-L), n = 58 axons (CC), n = 64 axons (IZ-contralateral). ****p <0.0001, ***p <0.001, **p <0.01, *p <0.05, ns, not significant by Kruskal-Wallis one-way ANOVA followed by two-tailed Dunn’s multiple comparisons test. Values are represented as a scatter plot with the mean indicated. f. Quantification of axon pausing events per hour in different cortical regions. N = 3 independent experiments, n = 45 axons (IZ-U), n = 35 axons (IZ-L), n = 40 axons (CC), n = 42 axons (IZ-contralateral). ****p <0.0001, ns, not significant by Kruskal-Wallis one-way ANOVA followed by two-tailed Dunn’s multiple comparisons test. Values are represented as a scatter plot with the mean indicated.

To study axon growth of CPNs, we electroporated neuronal progenitors *in utero* with a plasmid encoding the membrane-targeting motif of Lyn kinase (Lyn) fused with a fluorescent protein (FP) to visualise the overall neuron morphology at embryonic day E14.5 or E15.5^22–24^, followed by coronal brain sectioning 1–3 days after electroporation (**Fig. 1b**). Because average growth speed alone cannot distinguish continuous extension from intermittent growth interrupted by pauses, we quantified both axon growth speed and pausing events. We therefore performed long-term *in situ* imaging for up to 20 h, allowing us to compare these growth dynamics across different regions of the axonal trajectory.

After their initial formation in the upper intermediate zone (IZ-U), CPN axons turn and bundle with other axons in the lower intermediate zone (IZ-L) and head towards the corpus callosum (CC), before exiting the CC into the contralateral IZ^1,2,25^ (**Fig. 1a,c**). We found that in our slice culture, axons grew an order of magnitude faster than on 2D substrates. Growth speed was consistently above 60 µm/h, with notable differences among the regions. Axons in the IZ-U grew at a speed of 68.2 µm/h ± 16.4 µm/h (**Fig. 1d,e and Supplementary Video 1**) and paused 6.8 ± 1.8 times per h (**Fig. 1d,f and Supplementary Video 1**). Bundled axons in the IZ-L and the CC increased their growth speed by more than 20% (82.1 µm/h ± 22.3 µm/h and 88.1 µm/h ± 26.6 µm/h, respectively) (**Fig. 1d and Supplementary Video 1**) and reduced their pausing frequency to 4.9 ± 0.8 events per h and 5.1 ± 0.8 events per h, respectively (**Fig. 1d,f and Supplementary Video 1**). Upon exiting the CC and entering the contralateral IZ, axon growth speed decreased to 66.3 µm/h ± 22.8 µm/h, reaching velocities similar to those found in the ipsilateral IZ-U (**Fig. 1d,e**); in addition, the number of axon growth pauses increased to about 7.6 ± 1.3 pauses per h, reaching a pausing frequency similar to that in the ipsilateral IZ-U (**Fig. 1d,f**). Thus, growth speeds *in situ* were 12–21 times faster across the different regions than on 2D substrates and 2–4 times faster than in 3D^7^.

### Growth cones *in situ* recapitulate *in vivo* morphology and dynamics

Growth cones are thought to contain the machinery that drives axon growth^26,27^. Previous *in vivo* studies have described diverse growth cone morphologies during axon navigation, including fan-like and more streamlined forms^28–30^. We therefore asked whether the region-specific differences in axon growth dynamics are reflected at the level of the CPN growth cone. To address this, we performed *in situ* imaging of growth cone morphology and dynamics in Lyn-FP-labelled CPNs at defined positions along their growth trajectory.

Indeed, two growth cone morphologies dominated in specific regions: fan-like and spike-like (**Fig. 2a**). In the upper intermediate zone (IZ-U), where axons grew more slowly and paused more frequently, 79% of growth cones displayed a fan-like morphology, compared to only 28% and 10% in the IZ-L and CC, respectively) (**Fig. 2a-d**). In contrast, growth cones in the lower IZ (IZ-L) and the CC, where axons grew faster with fewer pauses, displayed a spike-like morphology (IZ-L: 72% and CC: 90%), compared to only 21% in the IZ-U (**Fig. 2a-d**). Moreover, we performed long-term live imaging to capture growth cone morphology and dynamics along the trajectories of individual axons *in situ*. In the IZ-U, growth cones predominantly displayed a fan-like morphology, which progressively decreased in size upon entering the IZ-L, adopting a spike-like morphology (**Extended Data Fig. 1a and Supplementary Video 2**). Conversely, spike-like growth cones in the contralateral IZ increased their size, adopting a fan-like morphology (**Extended Data Fig. 1b and Supplementary Video 2**). Axons therefore undergo a reversible morphological transition as they traverse distinct tissue regions. Since morphological changes were pronounced at the boundaries between regions, where axons change direction, we asked whether axon turning influences axon growth speed. Strikingly, we found that axon turning at the IZ-U/IZ-L border did not reduce growth speed (**Extended Data Fig. 1c,d and Supplementary Video 3**), contrasting with the behaviour of pioneering axons found in insects, as well as retinal ganglion cell (RGC) axons at the optic chiasm, which can stall for hours before turning^31–33^.

**Fig. 2:**
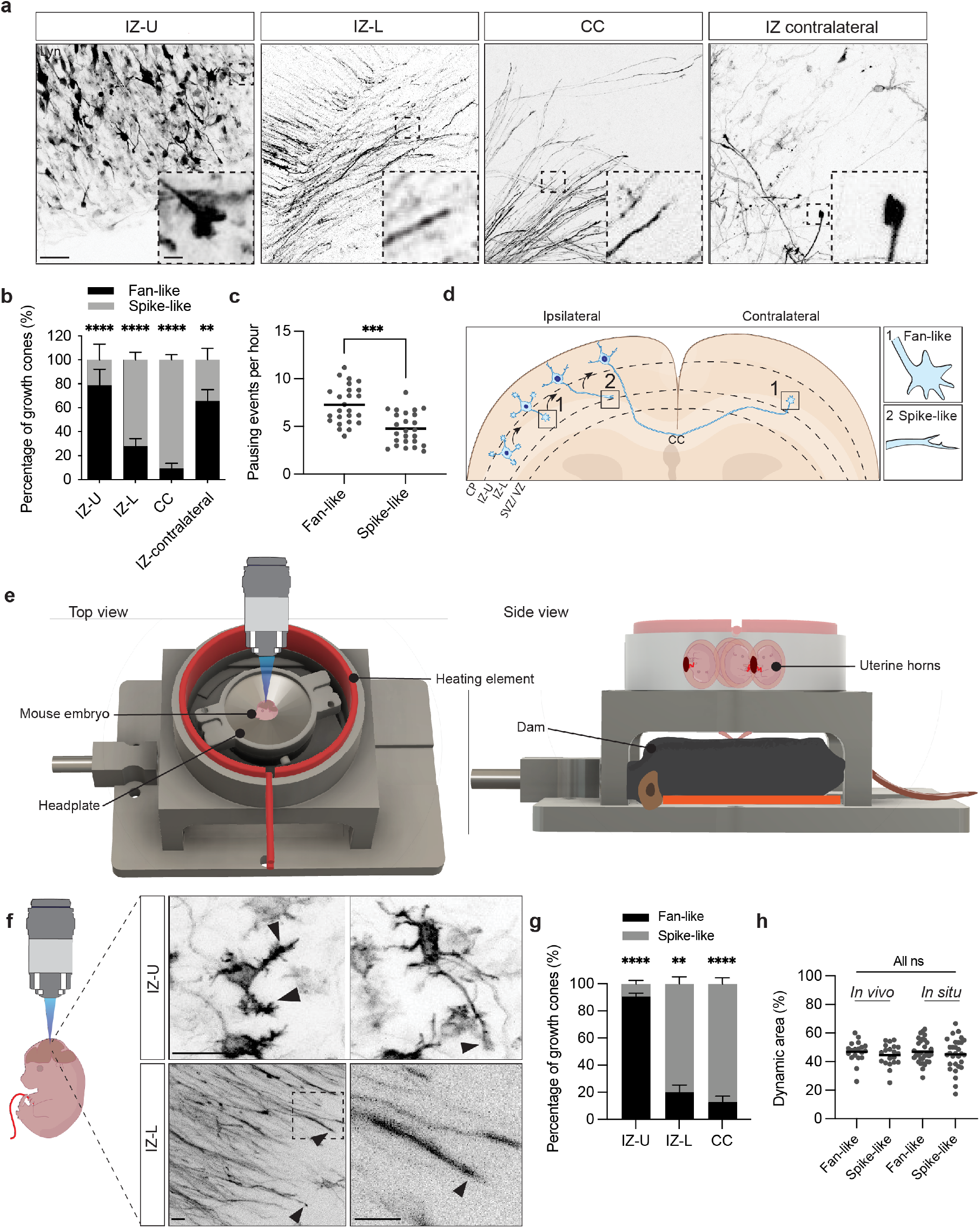
Region-specific growth cone morphologies *in situ* are conserved *in vivo*. a. Representative tiled images showing the location of each type of growth cone in the cortex. Neurons are labelled with Lyn-mNeonGreen via IUE. Insets highlight the different growth cone morphologies. Scale bar, 30 µm; inset scale bar, 5 µm. b. Percentage of fan-like and spike-like growth cones found in the different cortical regions analysed *in situ*. N = 3 independent experiments. n = 66 growth cones (IZ-U), n = 70 growth cones (IZ-L), n = 85 growth cones (CC), n = 57 growth cones (IZ-contralateral). ****p <0.0001, **p <0.01 by two-tailed Fisher’s exact test with Bonferroni correction for multiple comparisons (adjusted significance threshold α = 0.017). Each category (IZ-U, IZ-L, CC, IZ-contralateral) was compared independently between fan-like and spike-like using 2 × 2 contingency tables. Values are represented as stacked bars with standard deviation (SD) indicated. c. Quantification of the number of pausing events per hour for each growth cone type. N = 3 independent experiments, n = 25 fan-like growth cones, n = 25 spike-like growth cones. ***p <0.001by two-tailed Mann-Whitney U test. Values are represented as a scatter plot with the mean indicated. d. (Left) A schematic of growth cone morphologies at different regions of the cortex. (Right) Insets highlighting the different growth cone morphologies. e. A schematic showing top and side views of the *in vivo* embryo imaging platform AxoVivo. f. Representative images of growth cones in the IZ-U and IZ-L. Arrowheads mark the growth cones. Neurons are labelled with Lyn-mScarlet via IUE. Scale bars, 20 µm. g. Percentage of fan-like and spike-like growth cones found in the different regions analysed *in vivo*. N = 5 independent experiments. n = 18 growth cones (IZ-U), n = 29 growth cones (IZ-L), n = 27 growth cones (CC). ****p <0.0001, **p <0.01 by two-tailed Fisher’s exact test with Bonferroni correction for multiple comparisons (adjusted significance threshold α = 0.017). Each category (IZ-U, IZ-L, CC) was compared independently between fan-like and spike-like using 2 × 2 contingency tables. Values are represented as stacked bars with SD indicated. h. Quantification of the percentage of the relative dynamic area in fan-like and spike-like growth cones, *in situ* and *in vivo*. N = 4 independent experiments *in vivo*. n = 19 fan-like growth cones, n = 20 spike-like growth cones. N = 3 independent experiments *in situ*. n = 31 fan-like growth cones, n = 30 spike-like growth cones. ns, not significant by one-way ANOVA followed by two-tailed Tukey’s multiple comparisons test. Values are represented as a scatter plot with the mean indicated.

Because slice preparation disrupts the native tissue context, we asked whether these growth cone morphologies reflect genuine *in vivo* behaviour. Although recent advances in intravital imaging have enabled visualisation of neuronal activity, neuronal migration, and cell-cell interactions in the developing mouse brain^34–36^, resolving individual growing axons and their subcellular growth cone dynamics in the intact embryonic brain remains technically challenging. Building on our previous work using two-photon (2P) imaging to resolve axon and growth cone dynamics in regenerating axons in the spinal cord of adult mice^37–40^, we developed AxoVivo, an intravital imaging platform optimised to resolve subcellular structures at high spatial and temporal resolution while maintaining the embryo’s connection to the dam (**Fig. 2e**). Specifically, we constructed a multilayer setup, designed to maximise embryo stability by separating the embryo from the dam, maintaining the umbilical cord connection (**Fig. 2e and Extended Data Fig. 2 and Supplementary Video 4)**. Cortical blood flow remained undisturbed for over 6 h, supporting preservation of embryo viability and umbilical circulation (**Supplementary Video 5**).

2P imaging through the skull revealed that *in vivo* growth cone morphology was indistinguishable from that *in situ*: 91% of growth cones displayed a fan-like morphology in the IZ-U, whereas in the IZ-L and CC, growth cones mainly had a spike-like morphology (IZ-L: 80% and CC: 87%) (**Fig. 2f,g**). Moreover, quantification of dynamic growth cone areas showed that fan-like and spike-like growth cones displayed comparable relative dynamic areas *in vivo* and *in situ* (**Fig. 2h and Supplementary Video 6**), despite their distinct morphology. These values were similar to those previously reported for CNS axons growing in unconstrained 3D collagen matrices, but markedly higher than reported for growth cones on 2D substrates^7^. These similarities suggest that growth cone dynamics do not result from features disrupted by slice preparations.

AxoVivo thus provides direct optical access to axons and their growth cones in the living mammalian embryonic brain, bridging the gap between mechanically accessible brain section preparation and the physiological *in vivo* environment. Moreover, our data suggest that growth cones undergo a reversible morphological transition along their trajectory. Yet despite their distinct shapes, fan-like and spike-like growth cones showed comparable growth dynamics, indicating that growth cones of different morphologies share a common mode of advancing. This raised a central question: how do growth cones move forward through the tissue?

### Spike-like growth cones show reduced actin filament density in their central domain, whereas microtubules reach close to their leading edge

The cytoskeleton of the growth cone is thought to regulate axon growth and growth cone morphology^27^. We therefore sought to assess cytoskeletal dynamics *in situ*. Studying the cytoskeletal dynamics of growing axons *in situ* has been difficult as growth cones are small, highly dynamic structures and are located within dense, light-scattering tissue^1^. To overcome this, we built on our previously established slice imaging pipeline^22^ with targeted optimisation. Specifically, transferring slices to glass-bottom dishes immediately before imaging minimised the working distance between the objective and the tissue, enabling the use of high numerical aperture objectives that better resolve subcellular structures. We electroporated plasmids expressing the cytoskeletal markers Lifeact and the end-binding protein 3 (EB3), each fused to FPs to visualise actin filaments and polymerising ends of microtubules, respectively, in CPN growth cones, together with Lyn-FP^41,42^ (**Fig. 3**).

**Fig. 3:**
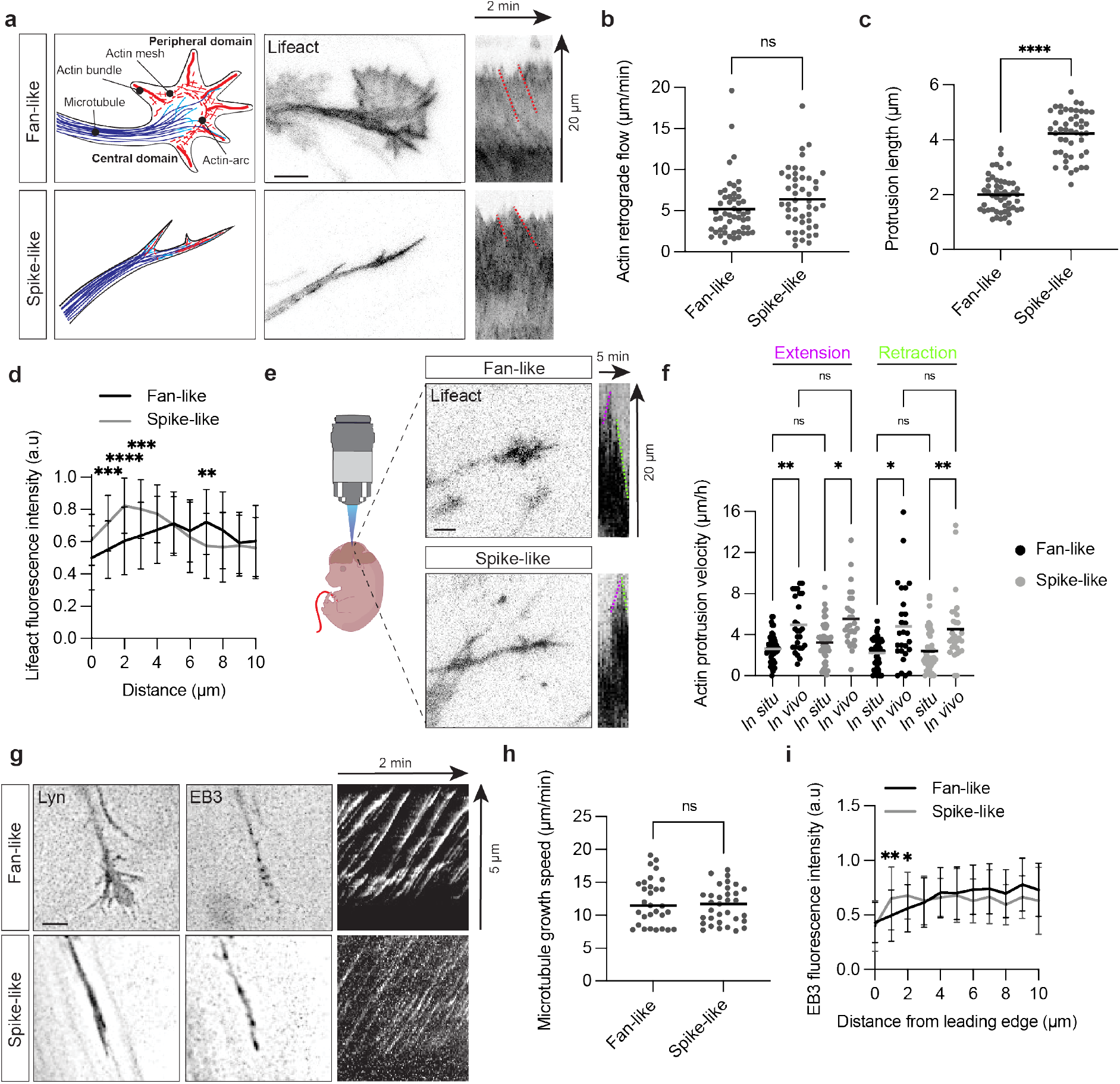
Cytoskeletal architecture underlies distinct growth cone morphologies. a. A schematic of the cytoskeleton of growth cones observed *in situ* (left). Representative images of the actin cytoskeleton in fan-like and spike-like growth cones *in situ* labelled with Lifeact-mNeonGreen via IUE (middle). Kymographs generated from the Lifeact-labelled growth cones (right). Red dashed lines in the kymograph indicate the slopes used to calculate actin retrograde flow speed in (b). Scale bar, 5 µm. b. Quantification of the speed of actin retrograde flow in each type of growth cone. N = 4 independent experiments, n = 51 fan-like growth cones, n = 47 spike-like growth cones. ns, not significant by two-tailed Mann-Whitney U test. Values are represented as a scatter plot with the mean indicated. c. Quantification of protrusion length in each growth cone type. N = 3 independent experiments, n = 51 fan-like growth cones, n = 47 spike-like growth cones. ****p <0.0001 by two-tailed Mann-Whitney U test. Values are represented as a scatter plot with the mean indicated. d. Quantification of actin fluorescence intensity in the growth cone. N = 3 independent experiments, n = 51 fan-like growth cones, n = 47 spike-like growth cones. Values are represented as line plots showing average fluorescence intensity of Lifeact ± SD. ****p = <0.0001, ***p <0.001, **p <0.01 by two-way ANOVA followed by two-tailed Šídák’s multiple comparisons test. e. Representative images of the actin cytoskeleton in fan-like and spike-like growth cones *in vivo*. Neurons are labelled with Lifeact via IUE. Kymographs were generated from the Lifeact-labelled growth cones (right). Examples of extension and retraction events quantified in (f) are marked on the kymographs with either magenta or green dashed lines, respectively. Scale bar, 10 µm. f. Quantification of actin protrusion extension and retraction speeds of each type of growth cone *in situ* and *in vivo*. N = 4 independent *in vivo* experiments, n = 27 fan-like growth cones, n = 28 spike-like growth cones. N = 3 independent *in situ* experiments, n = 51 fan-like growth cones, n = 47 spike-like growth cones. **p <0.01, *p <0.05, ns, not significant by Kruskal–Wallis one-way ANOVA followed by two-tailed Dunn’s multiple comparisons test. Values are represented as a scatter plot with the mean indicated. g. Representative images of fan-like and spike-like growth cones *in situ.* Neurons are labelled with Lyn-mScarlet (left) and end-binding protein 3 (EB3)-mNeonGreen (middle) via IUE, and kymographs are generated from the EB3-mNeonGreen-labelled growth cones (right). Scale bar, 5 µm. h. Quantification of microtubule growth speed. N = 3 independent experiments, n = 30 fan-like growth cones, n = 33 spike-like growth cones. ns, not significant by two-tailed Welch’s unpaired t-test. Values are represented as a scatter plot with the mean indicated. i. Quantification of EB3 fluorescence intensity in the growth cone. N = 3 independent experiments, n = 30 fan-like growth cones, n = 33 spike-like growth cones. Values are represented as line charts of the average mNeonGreen fluorescence intensity of EB3 ± SD. **p <0.01, *p <0.05 by two-way ANOVA followed by two-tailed Šídák’s multiple comparisons test.

Kymographic analysis revealed comparable rates of actin retrograde flow in fan-like and spike-like growth cones (**Fig. 3a,b and Supplementary Video 7**). The measured rates were similar to those previously reported in 3D collagen matrices^7^. Actin organisation differed between the two morphologies. Fan-like growth cones displayed shorter actin protrusions and enriched actin in the central (C) domain (**Fig. 3a,c,d**), whereas spike-like growth cones showed longer actin protrusions and increased distal actin in the peripheral (P) domain (**Fig. 3a,c,d)**.

We next asked whether these actin dynamics are preserved in the developing brain *in vivo*. We sought to exploit the potential of AxoVivo to resolve these subcellular dynamics. To this end, we used a mouse line with conditional expression of a Lifeact-mScarlet fusion protein^43^. Specifically, to follow actin dynamics in individual growth cones in the cortex of living embryos, we *in utero* electroporated a plasmid encoding Cre recombinase driven by Tubulin-α-1 (Tα1) to ensure neuron-specific Lifeact-mScarlet expression. Two days later, we imaged living embryos using our AxoVivo platform (**Fig. 3e**). While actin protrusion velocities were higher *in vivo* than *in situ*, we found no differences in the velocities between the growth cone morphologies (**Fig. 3f**).

We next asked whether microtubule behaviour could account for the differences in growth dynamics. Microtubule growth speeds, measured by EB3 comet dynamics, were also similar between growth cone morphologies (**Fig. 3g,h and Supplementary Video 8**). However, we found that microtubule plus ends protruded closer to the leading edge in spike-like growth cones compared with fan-like growth cones (**Fig. 3g,i**). Together, these data reveal that the two growth cone morphologies are distinguished primarily by cytoskeletal architecture rather than turnover rates, with reduced central actin in spike-like growth cones likely permitting microtubule plus ends to protrude closer to the leading edge. This raises the possibility that microtubules, rather than actomyosin, are the primary driver of axon growth in native CNS tissue.

### Microtubules are the primary driver of axon growth *in situ*

To test whether microtubules are the primary driver of axon growth *in situ*, we treated brain slices with the microtubule-depolymerising drug nocodazole, a treatment known to reduce axon formation and growth in cultured neurons^44–46^. Indeed, neurons exposed to nocodazole failed to form an axon *in situ*. Instead, they either retained a multipolar morphology or lacked discernible neurites (**Extended Data Fig. 3a,b**). Importantly, neurons with an axon already established before nocodazole treatment lost their polarised morphology and reverted to a multipolar or neurite-deficient state (**Fig. 4a,b and Supplementary Video 9**).

**Fig. 4:**
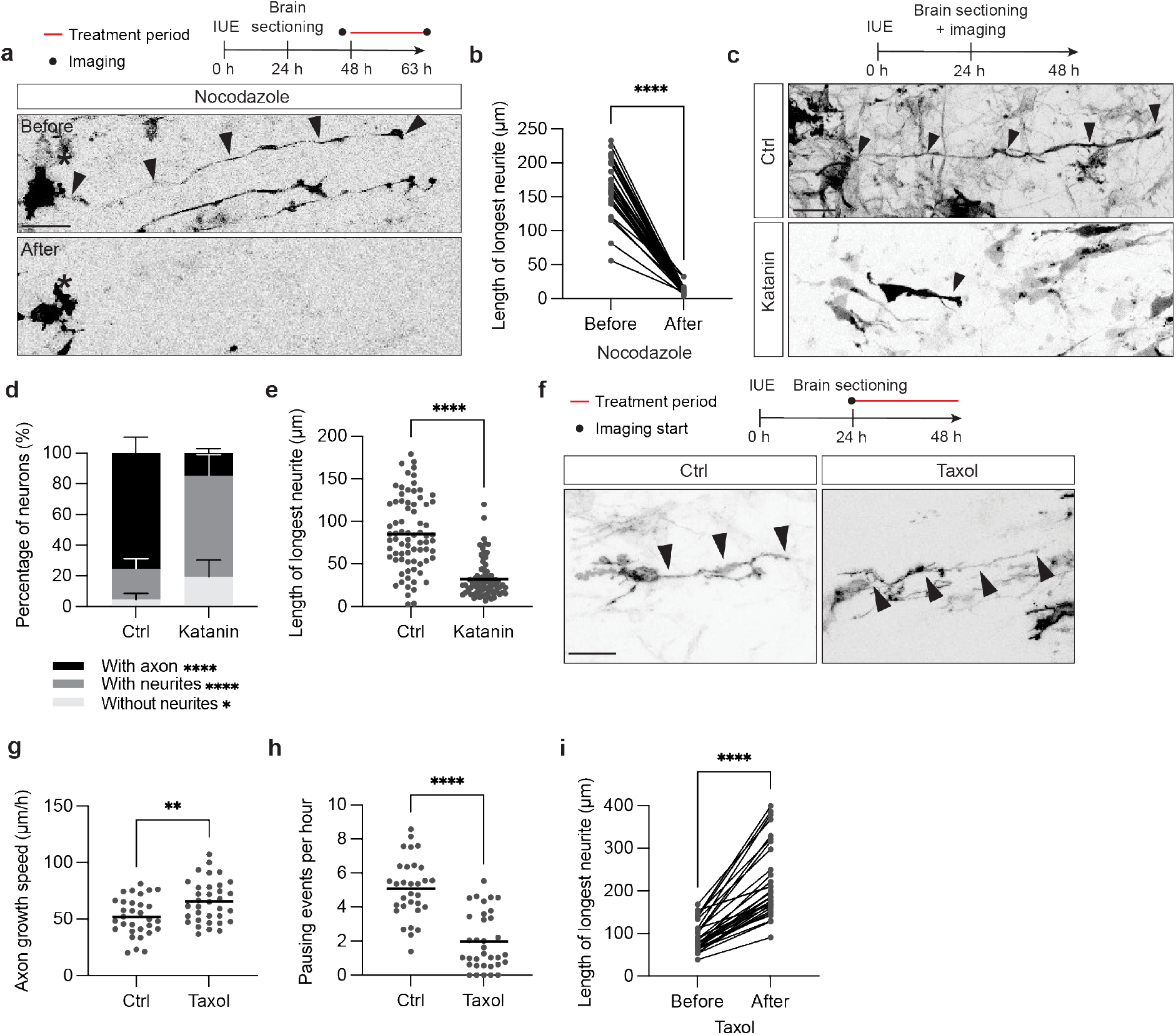
Axon growth relies on microtubule stabilisation *in situ*. a. Top, experimental design. Bottom, representative tiled images of a polarised neuron *in situ* before and 15 h after nocodazole treatment. Arrowheads label the axon before treatment. Asterisks indicate the initial position of the soma. Neurons are labelled with Lyn-mNeonGreen via IUE. Scale bar, 20 µm. b. Length of the longest neurite before and 15 h after nocodazole treatment. Each point corresponds to one neurite, with lines indicating paired observations. N = 3 independent experiments. n = 32 neurites. ****p <0.0001 by two-tailed Wilcoxon matched-pairs signed-rank test. Values are represented as a scatter plot. c. Top, experimental design. Bottom, representative images from tiled time-lapses of neurons expressing either katanin-mNeonGreen or only the empty plasmid (control) containing mNeonGreen via IUE. Arrowheads label the longest neurite. Scale bar, 20 µm. d. Percentage of neurons with an axon, with neurites or without neurites. N = 3 independent experiments. n = 78 neurons (control), n = 90 neurons (katanin). ****p <0.0001, *p <0.05 0.0181 by two-tailed Fisher’s exact test with Bonferroni correction for multiple comparisons (adjusted significance threshold α = 0.017). Each cell category (with axon, with neurite, without neurite) was compared independently between control and katanin using 2 × 2 contingency tables. Values are represented as stacked bars with SD indicated. e. Length of the longest neurite in control and katanin-overexpressing neurons in (c). N = 3 independent experiments. n = 75 neurons (control), n = 74 neurons (katanin). ****p <0.0001 by two-tailed Mann-Whitney test. Values are represented as a scatter plot with the mean indicated. f. Top, experimental design. Bottom, representative images from tiled time-lapses of either control or taxol-treated neurons *in situ*. Arrowheads label the longest neurite. Neurons are labelled with Lyn-mNeonGreen via IUE. Scale bar, 20 µm. g. Quantification of axon growth speed in the IZ-U. N = 3 independent experiments, n = 31 axons (control), n = 34 axons (taxol). **p <0.01by two-tailed Mann-Whitney test. Values are represented as a scatter plot with the mean indicated. h. Quantification of axon pausing events per hour in the IZ-U. N = 3 independent experiments, n = 31 axons (control), n = 34 axons (taxol). ****p <0.0001 by two-tailed Mann-Whitney test. Values are represented as a scatter plot with the mean indicated. i. Length of the longest neurite before and 15 h following taxol treatment. N = 3 independent experiments. n = 31 neurites. ****p <0.0001 by two-tailed Wilcoxon matched-pairs signed-rank test. Values are represented as a scatter plot.

As nocodazole acts on neurons and non-neuronal cells within the slices, we next used a cell-autonomous approach to depolymerise microtubules. To this end, we generated a conditional expression plasmid enabling neuron-specific overexpression of the microtubule-severing protein katanin^47^ together with a fluorescent reporter under a Tα1-driven Cre cassette^48^. Under these conditions, neurons failed to form an axon (**Fig. 4c-e and Supplementary Video 10**). Moreover, cell-autonomous expression of spastin, another microtubule-severing protein, resulted in a similar lack of axon formation (**Extended Data Fig. 3c-e and Supplementary Video 11**).

To directly assess whether increasing microtubule protrusion is sufficient to promote axon growth, we stabilised microtubules pharmacologically with low doses of taxol. Indeed, this caused a significant increase in axon growth speed *in situ* (**Fig. 4f,g**). We found a 26% increase in axon growth speed and a 61% decrease in axon pausing following taxol application (**Fig. 4g,h and Supplementary Video 12**), as well as an increase in longest-neurite length 15 h after taxol treatment (**Fig. 4i**). Taken together, these results show that microtubules are the primary driver of axon growth *in situ*.

### Myosin II regulates soma migration and growth cone shape but not axon growth

If microtubules are the primary driver of axon growth in its natural habitat, this raises the possibility that actomyosin-mediated pulling via adhesion sites plays a less important role in this process. Therefore, if axons can extend *in situ* without pulling on the ECM, they should be able to grow independently of the actin motor myosin II, which is necessary for adhesion-dependent movement^49,50^. To test this possibility, we treated neurons in brain slices with the pharmacological myosin II inhibitor blebbistatin^49,51,52^. Indeed, we found that blebbistatin did not affect growth speed in either the IZ or the CC (**Fig. 5a-c** **and Supplementary Video 13**). Interestingly, however, blebbistatin application changed axonal growth cone morphology in the IZ-U, from around 20% to more than 80% of growth cones in the IZ-U exhibiting a spike-like morphology (**Fig. 5d**). Moreover, we found that blebbistatin reduced the movement of the radially migrating CPN cell bodies (**Fig. 5e,f**). Thus, myosin II regulates growth cone morphology and CPN soma migration, but not axon growth *in situ*.

**Fig. 5:**
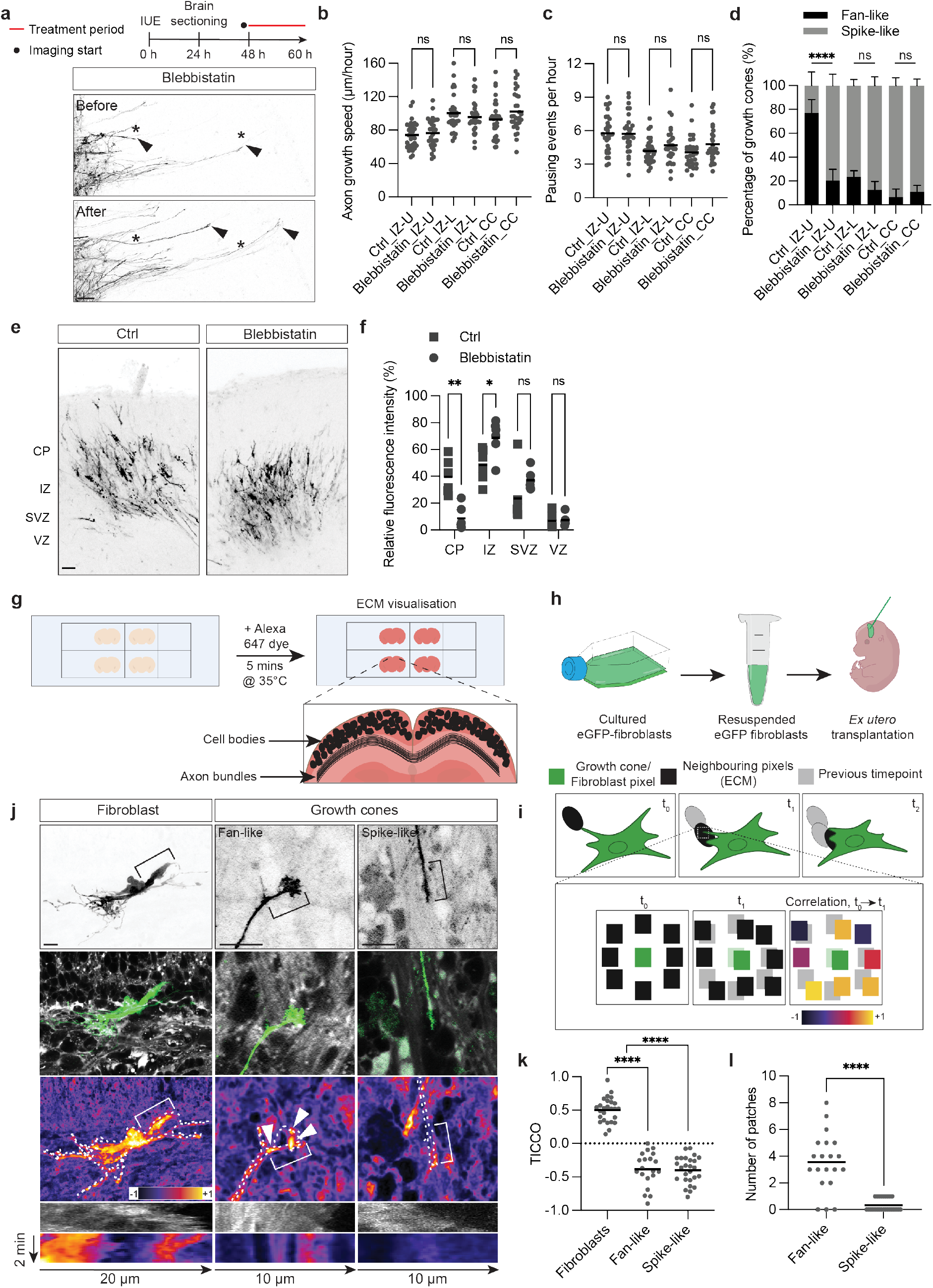
Growth cones lack mechanical coupling to the ECM. a. Top, experimental design. Bottom, representative tiled images of axon growth in the IZ-L before and 10 h after blebbistatin application. Arrowheads indicate the tip of growing axons. Asterisks mark the initial position of the axon tip. Neurons are labelled with Lyn-mNeonGreen via IUE. Scale bar, 50 µm. b. Quantification of axon growth speed in the IZ-U, IZ-L and CC, under control or blebbistatin-treated conditions. N = 3 independent experiments, n = 60 axons (IZ-U), n = 60 axons (IZ-L), n = 60 axons (CC). Data analysed as control vs blebbistatin: ns, not significant by two-tailed Welch’s unpaired t-test. Values are represented as a scatter plot with the mean indicated. c. Quantification of axon pausing events per hour in the IZ-U, IZ-L and CC, either control or blebbistatin-treated. N = 3 independent experiments, n = 60 axons (IZ-U), n = 60 axons (IZ-L), n = 60 axons (CC). ns, not significant by two-tailed Welch’s unpaired t-test. Values are represented as a scatter plot with the mean indicated. d. Quantification of the percentage of fan-like and spike-like growth cones in the IZ-U, IZ-L and CC, either control or blebbistatin-treated. N = 3 independent experiments, n = 60 axons (IZ-U), n = 60 axons (IZ-L), n = 60 axons (CC). ****p <0.0001, ns, not significant by two-tailed Fisher’s exact test with Bonferroni correction for multiple comparisons (adjusted significance threshold α = 0.017). Values are represented as stacked bars with SD indicated. e. Representative tiled images of neurons in control (left) and blebbistatin-treated (right) brain slices. Neurons are labelled with Lyn-mNeonGreen via IUE. Scale bar, 50 µm. f. Quantification of soma migration data in (e). n = 6 electroporated brains. Data analysed as control vs blebbistatin. **p <0.01; *p <0.05; ns, not significant by two-tailed Welch’s unpaired t-test. Values are represented as a scatter plot with the mean indicated. g. A schematic of the protocol used for brain section labelling for Super-Resolution Shadow Imaging (SUSHI). h. A schematic of the protocol used for *ex utero* eGFP-fibroblast transplantation into the cortex. i. A schematic of the time-integrated cross-correlation (TICCO) pipeline used to analyse fibroblast-extracellular matrix (ECM) or growth cone-ECM interaction. j. Representative images of a fibroblast and growth cones imaged using SUSHI (top and middle rows) and heatmaps generated via TICCO analysis (bottom row). Manual tracings of the fibroblast and growth cones are overlaid on the heatmaps (white dotted tracings). Arrowheads mark patches in the fan-like growth cone. Square brackets label the areas from which the kymographs below were generated, either from the fibroblast/growth cone (upper kymograph) or from the TICCO heatmap (lower kymograph). Scale bar, 10 µm. k. Quantification of fibroblast-ECM or growth cone-ECM movement correlation, analysed using TICCO. N = 3 independent experiments, n = 26 fibroblasts, n = 20 growth cones (fan-like), n = 26 growth cones (spike-like). ****p <0.0001 by one-way ANOVA followed by two-tailed Tukey’s multiple comparisons test. Values are represented as a scatter plot with the mean indicated. l. Quantification of the number of patches observed in the growth cones analysed in (j). N = 3 independent experiments, n = 20 growth cones (fan-like), n = 26 growth cones (spike-like). ****p <0.0001 by two-tailed Mann-Whitney U test. Values are represented as a scatter plot with the mean indicated.

### Axons grow without pulling on the ECM

As axons grow *in situ* without myosin II activity, it raises the possibility that axon growth and growth cone advancement can occur without pulling on their native ECM. We therefore sought to measure growth cone–ECM pulling. Standard approaches for quantifying mechanical cell– ECM coupling, such as particle image velocimetry or kymograph analysis, work best in simplified environments with limited background motion^53,54^. In the native developing brain, however, the ECM undergoes continuous structural remodelling^4^, generating background displacement signals that make it difficult to distinguish displacement caused by cellular pulling. Existing methods therefore cannot reliably detect whether a moving cell is exerting a pulling force on its surroundings in this physiological setting. To overcome this limitation and directly assess mechanical coupling between growth cones and the ECM *in situ*, we developed a pipeline to observe correlated movement, in which we co-imaged Lyn-FP labelled growth cones of CPNs *in situ* together with the surrounding ECM using SUSHI^4^ (**Fig. 5g**). As a positive control for detecting correlated movement, we exogenously added fibroblasts, which mechanically couple to their environment to migrate^55^, to the cortex before sectioning (**Fig. 5h**). To quantify this cellular behaviour in relation to the ECM, we developed a four-dimensional (three spatial dimensions and time) time-integrated cross-correlation (TICCO) analysis tool whereby the movement of the labelled cells was correlated to the surrounding ECM at single-pixel resolution (**Fig. 5i**).

Indeed, we found that fibroblast movement positively correlated with the movement of the surrounding ECM (**Fig. 5j,k and Supplementary Video 14**). Importantly, this correlated movement depended on myosin II, as it was abrogated upon blebbistatin treatment (**Extended Data Fig. 4a,b**). As an internal physiological positive control, we used the CPN soma, which migrates to the cortical plate along radial glia through adhesion sites, while the axon grows from its trailing edge^48,56–58^. Similarly, we found that the migratory movement of the CPN soma positively correlated with the movement of the surrounding ECM (**Extended Data Fig. 4c,d**).

Importantly, when we used our TICCO analysis pipeline to detect potential coupling between growth cones and the surrounding ECM, we found such coupling to be largely undetectable (**Fig. 5j,k and Supplementary Video 14**). Only in fan-like growth cones did we occasionally observe a small number of dynamic patches exhibiting high local correlation with the ECM (**Fig. 5j,l and Supplementary Video 14**). Together, these results indicate that axonal growth cones *in situ* extend without exerting measurable pulling on the ECM. Thus, cortical axons grow in their native environment with a low-adhesion, microtubule-driven mechanism.

## Discussion

How do axons achieve the rapid growth necessary to wire the mammalian brain? Our findings revealed that mammalian CPN axons grow in their native environment without pulling on the surrounding ECM, thereby employing a mode of growth that is both faster and mechanically simpler than adhesion-dependent traction. AxoVivo enabled us to test whether the behaviours identified in brain slices are preserved in the living embryonic brain. By combining physiological context with subcellular imaging access, it bridges the gap between slice preparation and the intact mammalian embryo. The close correspondence between *in situ* and *in vivo* growth cone behaviour supports the physiological relevance of the mechanisms dissected in slices.

The developing cortex exposes axons to changing cellular densities, extracellular organisation, guidance landscapes, and mechanical properties^18,59–62^. Growth cone morphology switched markedly between fan-like and spike-like across regions; changes in growth speed were relatively modest, within approximately 20%. This contrasts with the drastic halting of axon growth observed at guidance choice points in other systems: grasshopper Ti1 pioneer axons stall for many hours at limb segment boundaries and expand large fan-like growth cones before turning^28,31,32^, and zebrafish motor axons in the spinal cord pause for 1–2 hours at intermediate targets before resuming growth^63^. This suggests that mammalian CPN axons in their native environment are capable of maintaining growth while turning and rearranging their trajectory. Hence, growth cone morphology and axon growth appear to be mediated through different molecular events.

Indeed, myosin II inhibition converted fan-like growth cones to a spike-like morphology without affecting axon growth speed, directly dissociating growth cone morphology from growth speed. This suggests that in the mammalian CNS, growth cone morphology and axon growth speed are not necessarily causally coupled, and that the actin cytoskeleton acts as a regulator of growth cone architecture rather than a driver of axon elongation. Moreover, the actin cytoskeleton plays a more dominant role in many developmental processes, including asymmetric cell division^64^, branching^65,66^ and scaffold formation during synapse establishment^67,68^. This is further supported by our recent finding that myosin II together with Arp2/3 forms an oscillatory network during initial neuronal polarisation, which is then lifted once the axon is specified and continues to grow^44^. Moreover, microtubules have been shown to interact with the plasma membrane and cell cortex during protrusion, either by generating local pushing forces or by regulating membrane-proximal actin assembly^69–73^. Our data suggest a model in which growth cone microtubules drive forward protrusion in native tissue.

Several studies have established that migrating non-neuronal cells move fastest in the absence of stable adhesions^9,74^. Building adhesion sites to enable cellular traction and their subsequent disassembly at the cell rear to release forward movement are time-consuming processes that constrain mesenchymal migration to comparatively slow speeds^11,13^. Consistent with this principle, CPN axons grew rapidly *in situ* without exerting detectable pulling on the surrounding densely confined ECM^75,76^. In such densely confined spaces, non-neuronal cells can migrate faster without adhesion by exploiting friction against confining walls, a mechanism conceptually analogous to chimney climbing^77^. Moreover, and consistent with mesenchymal migration being an inherently slower motility mode, CPN soma migration proceeds at least an order of magnitude slower than axon growth *in situ*^48^, placing it firmly in the range of adhesion-dependent cell migration rather than the rapid protrusion-driven advance we describe here.

This dissociation between axon growth and soma migration is remarkable given the distinctive mechanical challenge CPNs face during corticogenesis. While the soma migrates radially toward the cortical plate along glial fibres, a process requiring actomyosin-dependent force generation and mechanically coupled movement, the axon extends simultaneously along an orthogonal trajectory^56,78–80^. Our findings suggest that CPNs resolve this challenge by operating two distinct motility programmes within the same cell: a mechanically coupled, actomyosin-dependent programme for soma migration, and a mechanically uncoupled, microtubule-driven programme for axon growth^48,74^. This separation is likely functionally necessary. If axon growth relied on stable substrate adhesion, the growing axon shaft would risk becoming mechanically tethered to the radial glial scaffold used by the migrating soma^56,79^, potentially constraining the axon to follow the soma’s radial trajectory. A low-adhesion mode of growth allows the axon to navigate its own path independently, enabling long-range connectivity to be established while radial migration proceeds concurrently.

Our findings provide a new perspective on microtubule-related developmental disorders, such as Tubulinopathies, which are associated with cortical malformations, including abnormalities of the corpus callosum and axon growth defects^81–83^. From this perspective, defects in tubulin function may impair the microtubule-protrusion machinery that drives axon growth in the developing brain. Reactivating developmental growth programmes is a strategy for promoting regeneration in the injured adult CNS, including via several regeneration-promoting interventions that converge on cytoskeletal mechanisms^19,38,45,84–91^. Our results here suggest that promoting axon regeneration may require shifting growth cone behaviour toward the microtubule-driven fast-growing state we describe, thereby providing a more targeted strategy for stimulating axon regeneration in the adult CNS. Together, our findings identify a low-adhesion microtubule-based mechanism that axons employ in their native environment to grow rapidly, resulting in efficient brain wiring.

## Methods

### Data availability

All data reported in this paper and any additional information required to reanalyse the data will be shared upon request. Requests should be directed to and will be fulfilled by the lead contact, Frank Bradke.

### Materials availability

The plasmids generated in this study will be deposited in Addgene.

### Code availability

The TICCO Python script will be deposited on GitHub.

### Animals

All animal experiments were performed following the Animal Welfare Act and the guidelines of the North Rhine-Westphalia State Environment Agency (Landesamt für Verbraucherschutz und Ernährung (LAVE)). The animals were single-housed at a controlled room temperature of 21–22 °C, and an artificial 12 h light:dark cycle (lights off at 18:00). Mice were given food and water *ad libitum* throughout the experiment.

Timed-pregnant WT C57BL/6J mice (Janvier Labs), Actin-eGFP^92^ or Lifeact-mScarlet C57BL/6N^43^ mice were used for IUE. The embryos were allowed to develop to the indicated time points depending on the experiment. Embryonic sex was not determined in this study, resulting in a random mix of both sexes in all related experiments.

### Plasmids

For imaging axon and growth cone morphology and dynamics *in situ*, plasmids encoding the membrane-targeting motif of Lyn kinase fused to either mScarlet or mNeonGreen were used. The plasmid pCAG-loxSTOPlox-Lyn-mNeonGreen (Addgene plasmid #175257^22^) was used as a template to generate pCAG-loxSTOPlox-Lyn-mScarlet. Lyn-mNeonGreen was excised from pCAG-loxSTOPlox-Lyn-mNeonGreen using PstI/SacI. The Lyn mScarlet sequence was amplified by PCR from pCAG-lox-rox-STOP-rox-Lyn-mScarlet-lox-Lyn-mNeonGreen (Addgene plasmid #175259^22^) and cloned into the linearised pCAG-loxSTOPlox backbone using Gibson assembly (HiFi DNA Assembly Cloning Kit; New England Biolabs), yielding pCAG-loxSTOPlox-Lyn-mScarlet. For *in vivo* imaging of axon and growth cone dynamics, the fluorophore in the plasmid pCAG-loxSTOPlox-Lyn-mNeonGreen (Addgene plasmid #175257^22^) was replaced with mGreenLantern (a gift from Gregory Petsko^93^, Addgene plasmid #164461) using PCR. To image actin dynamics *in situ*, pTub-alpha1-Lifeact-mNeonGreen was used (Addgene plasmid #175258^22^). For imaging microtubule dynamics *in situ*, pTub-alpha1-EB3-mNeonGreen was used (Addgene plasmid #175263^22^). For the overexpression of the microtubule-severing proteins, katanin and spastin, the plasmids were generated as follows: The backbone, pCAG-loxSTOPlox-mNeonGreen, was amplified from the plasmid pCAG-loxSTOPlox-Lyn-mNeonGreen (Addgene plasmid #175257^22^) using PCR. Katanin or spastin inserts were also amplified by PCR from p80-Katanin-mNeonGreen (a gift from Daniel Gerlich^47^, Addgene plasmid #191762) and HA-eGFP-Spastin-M87-with-exon-4 (a gift from Tarun Kapoor^94^, Addgene plasmid #167542), respectively. The backbone and inserts were assembled using Gibson assembly (HiFi DNA Assembly Cloning Kit; New England Biolabs), yielding either pCAG-loxSTOPlox-katanin-mNeonGreen or pCAG-loxSTOPlox-spastin-mNeonGreen. All floxed plasmids were co-expressed with Tα1-Cre (Addgene plasmid #133924^89^), resulting in neuron-specific expression.

### *In utero* electroporation (IUE)

IUE was performed as previously described ^22^, with minor modifications. Briefly, E14.5 or E15.5 pregnant female WT C57BL/6J mice (Janvier) or Lifeact-mScarlet C57BL/6N ^43^ received a subcutaneous injection of buprenorphine (WDT; stock 0.03 mg/ml, diluted 1:10 in 50% glucose; 0.1 mg/kg mouse) 20 min before anaesthesia induction with isoflurane (Abbott; 5% for induction and 1.5–2% for maintenance). A small abdominal incision was made to expose the uterine horns, which were regularly moistened with warm saline to prevent dehydration and hypothermia.

Both lateral ventricles were injected with 2–3 µl of a plasmid mixed with 10% Fast Green dye (Sigma) using glass micropipettes with a pipette puller (Zeitz). Electroporation was performed using platinum tweezertrodes connected to an ECM 830 pulse generator (BTX Harvard Apparatus). Five square-wave pulses (35 V, 600 ms intervals, 50 ms duration) were delivered to target the somatosensory cortex.

Following electroporation, the uterine horns were returned to the abdominal cavity, and the incision was sutured and clipped. Mice were placed under a far-infrared warming lamp for 20 min before being returned to the holding room. The mice were additionally given high-calorie food (Ssniff) and sunflower seeds (Pauls Mühle). An additional dose of buprenorphine (0.1 mg/kg) was administered 4–6 h after IUE. Postoperative analgesia was further provided via tramadol in the drinking water (Venusberg Apotheke Bonn; stock 100 mg/ml, diluted 1:1000; final concentration 0.1 mg/ml) for up to 3 days after IUE.

### Brain sectioning

Brain sections were prepared as previously described ^22^ with slight modifications. Embryos were collected 1, 2 or 3 days after IUE. For embryo collection, pregnant mice received a subcutaneous injection of buprenorphine (WDT; stock 0.03 mg/ml, diluted 1:10 in 50% glucose, 0.1 mg/kg) 20 min prior to anaesthesia induction with isoflurane (Abbott; 5% for induction and 1.5–2% for maintenance). The abdominal incision was reopened, embryos were collected, and immediately sacrificed by decapitation. Embryonic heads were placed in ice-cold Hank’s Balanced Salt Solution (HBSS) supplemented with 0.5% glucose (Sigma). Following embryo collection, the pregnant mouse was euthanised by cervical dislocation.

Brains were dissected, embedded in 3% low-melting-point agarose (Biozyme), and sectioned into 150–180 µm coronal slices using a VT1200S vibratome (Leica). Slices were transferred onto 30 mm polytetrafluoroethylene membranes (Millipore) placed in 35 mm transwell dishes (Fluorodish, WPI). Slice culture medium consisting of Neurobasal (1×), Fetal calf serum (FCS; 5%), B27 supplement (1:50), GlutaMAX (1:400), penicillin/streptomycin (1:200) (all from Thermo Fisher Scientific), horse serum (5%), Neuropan-2 supplement (1:100) (both from Pan-Biotech; final pH = 7.3) was added beneath the membrane. Slices were incubated at 37°C and 5% CO_2_ for at least 1 h prior to imaging.

For drug treatments, blebbistatin (40 µM; B0560, Sigma-Aldrich), nocodazole (5 µM; M1404, Sigma-Aldrich) or taxol (6 nM; Cay10461-25, Biomol) was diluted in slice culture medium, and 500 µl was added to the dish beneath the slices before imaging. Control experiments received medium from the same batch without the drug.

### *Ex utero* fibroblast transplantation

Meningeal eGFP-expressing fibroblasts were isolated from Actin-eGFP mice. Brain meninges were collected in HBSS and then digested with trypsin/DNase solution at 37°C for 15 min. The trypsin/DNase solution was removed, and the tissue was washed with 1× HBSS supplemented with HEPES buffer. Fibroblasts were manually dissociated and cultured in Minimum Essential Medium (MEM) supplemented with 10% FCS. The medium was replaced within 24 h to remove cell debris.

For transplantation, fibroblasts were used after at least 3 passages. Shortly before transplantation, fibroblasts were trypsinised at 37°C for 5 min using 0.05% trypsin, which was subsequently inactivated with MEM containing 10% FCS. The cell suspension (30,000 to 50,000 cells) was prepared in GlutaMAX (Thermo Fisher Scientific), penicillin/streptomycin (Life Technologies), glucose, and 10% Fast Green dye (Sigma). A volume of 2–3 µl of the fibroblast suspension was injected bilaterally into the cortex. Following transplantation, embryos were collected in ice-cold HBSS supplemented with 0.5% glucose for at least 30 min prior to brain dissection.

### Confocal imaging

Long-term axon growth and growth cone imaging were performed using a confocal LSM880 microscope (Zeiss) equipped with an incubation chamber maintained at 35°C and 5% CO_2_. For long-term axon growth imaging, either a Plan-Apochromat 10×/0.45 M27-FWD = 2.1 mm dry objective (Zeiss) or a C-Achroplan 32×/0.85 W Corr M27 VIS-IR objective (Zeiss) was used. Images were acquired as tiled 100–180 µm z-stacks with a step size of 1–3 µm and collected every 5 min for 15–20 h. When tiling was applied, images were stitched using the ZEN Black software (Zeiss).

For the imaging of growth cone morphology and actin dynamics, the C-Achroplan 32×/0.85 W Corr M27 VIS-IR objective was used. Images were acquired as 5–15 µm z-stacks with a step size of 0.2 µm and collected every 2–3 s for 2–3 min.

To assess migration defects following blebbistatin treatment, slices were incubated with medium containing 40 µM blebbistatin for 24 h prior to imaging. Imaging was performed using the C-Achroplan 32×/0.85 W Corr M27 VIS-IR objective. Images were acquired as 50 µm z-stacks with a step size of 3 µm.

For SUSHI imaging, an LSM980 microscope (Zeiss) equipped with an incubation chamber maintained at 35°C and 5% CO2 was used. Images were acquired in Airyscan super-resolution mode. Slices were transferred to 8-well glass-bottom dishes (Ibidi) shortly before imaging, and 5 µl of pre-warmed 40 µM Alexa Fluor 647 dye (Thermo Fisher Scientific) was applied directly to each slice. Slices were incubated at 35°C for 5 min prior to imaging. Images were acquired as 20 µm z-stacks with a 0.5 µm step size using a Plan-Apochromat 40×/1.4 oil-immersion objective.

### Widefield imaging

Microtubule dynamics were imaged using a DeltaVision Elite system equipped with an incubation chamber maintained at 37°C and 5% CO_2_. A 60×/1.42. oil-immersion objective was used together with eGFP/mCherry or eGFP/TRITC dichroic mirrors and the corresponding emission and excitation filters. Images were acquired every 1–2 s for 2–3 min as 2 µm z-stacks with a step size of 0.2 µm. The time-lapse sequences were deconvolved using DeltaVision SoftWoRx software.

### AxoVivo platform

The custom-made AxoVivo platform was designed for high-resolution *in vivo* imaging and comprised three layers. The lowermost layer consisted of a microscope insert plate to which the remaining components were attached. The middle layer contained a bath for the mouse embryos, with a bottom opening through which the exteriorised uterine horns were inserted. The uppermost layer consisted of a rotating lid equipped with an adjustable iris (Thorlabs) to secure individual embryos and a headplate that could be screwed into place to further immobilise the embryo during imaging. The bath and isoflurane mask were 3D-printed in clear resin using a Form 3+ printer (Formlabs). All other components were printed from biocompatible polylactic acid (PLA) filament using an UltiMaker S7 printer (UltiMaker). The bath and isoflurane mask were secured with magnetic cubes to stabilise the setup during imaging. The bath temperature was maintained using a 12 V silicone heating strip.

### Embryo immobilisation for *in vivo* imaging

Two or three days following IUE, the mother was injected subcutaneously with buprenorphine (WDT, stock 0.03 mg/ml, diluted 1:10 in 50% glucose, 0.1 mg/kg mouse) 20 min before being anaesthetised with isoflurane (5% for induction and 1.5% for maintenance). The incision was carefully reopened. Both uterine horns were exteriorised and placed inside the AxoVivo bath that sat on top of the pregnant mouse. The hole in the holder was sealed using silicone adhesive (WPI). The bath was then filled with artificial cerebrospinal fluid (aCSF) that was maintained at 37 °C using a heating element. A small incision was then made in the uterine wall to expose the embryo, which was then immobilised using an adjustable iris (Thorlabs) and a custom-made headplate.

### Two-photon imaging

For imaging growth cone dynamics *in vivo*, an upright Bergamo II series multiphoton microscope (Thorlabs) equipped with an Apo LWD 25×/1.10 water immersion objective (Nikon) with a working distance of 2 mm was used. mGreenLantern was excited at 940 nm using the galvo-resonant scanner. Images were acquired as 10–50 µm z-stacks with a 1 µm step size. Images were acquired every 15–20 s, depending on z-stack size.

For imaging actin dynamics, an upright Axio Examiner LSM7MP multiphoton microscope (Zeiss) equipped with an InSight X3 tunable infrared laser (Spectra-Physics) and an Apo LWD 25×/1.10 water immersion objective (Nikon) with a working distance of 2 mm was used.

mScarlet was excited at 1050 nm. mScarlet emission was bandpass filtered (655/40) and detected by an ultra-sensitive non-de-scanned gallium-arsenide-phosphide (GaAsP) detector. Images were acquired as 10–50 µm z-stacks with a 1 µm step size. Time-lapse images were acquired as a single-plane series of 1024 × 1024 pixels with a 10 s interval.

For blood flow imaging, the same LSM7MP mentioned above was used. Time-lapse images were acquired as a single-plane series of 1024 × 1024 pixels with a 0.5 s interval in epi-fluorescence mode using an AxioCam (Zeiss) camera with an eGFP filter set and a HXP 120 mercury lamp (Zeiss) as a light source.

### Analysis

#### Morphology, speed and pausing

Axon growth speed was quantified by manual tracking using the “Spots” function in Imaris (v9.9). Spot diameter was set to 4.5 µm, and tracking was performed using the “Autoregressive motion” algorithm. An axon growth event of <0.5 µm over a 5-min interval was considered a pausing event. For axon growth speed tracking at turning points, the speed was measured from 12 frames (60 min) before the start of the turning event to 12 frames (60 min) after the end of the turning event.

Cortical regions were identified based on anatomical landmarks, cytoarchitecture, cell density, and axon tract organisation in coronal embryonic slices. The ventricular zone (VZ) was identified as the densely packed proliferative layer lining the lateral ventricle, whereas the subventricular zone (SVZ) was identified as the region immediately basal to the VZ. The intermediate zone (IZ) was identified as the cell-sparse region between the SVZ and cortical plate (CP), containing migrating neurons and extending axons. For regional analysis, the IZ was subdivided into an upper intermediate zone (IZ-U), located adjacent to the CP, and a lower intermediate zone (IZ-L), adjacent to the SVZ and enriched in bundled callosal axons. The CP was defined as the dense layer of post-mitotic neurons located superficial to the IZ and below the marginal zone. The corpus callosum (CC) was identified as the dense commissural axon tract crossing the midline between the two hemispheres.

Growth cone morphology was classified based on the presence or absence of lamellipodia. Growth cones containing lamellipodia were classified as fan-like, whereas growth cones lacking lamellipodia were classified as spike-like.

Neurons in slices treated with nocodazole and neurons overexpressing katanin and spastin were quantified as follows: Neurons were manually tracked using the “Filaments” tool in Imaris (v9.9) and considered to possess an axon if the longest process was ≥50 µm in length. Neurons lacking any process ≥5 µm were classified as having no neurites.

To quantify the active area of growth cones, a maximum intensity projection was generated from each image series using Fiji (v2.16.0)^95^. Highly dynamic regions undergo rapid local intensity changes and therefore appear brighter in the maximum projection compared to more static areas. The contour of each growth cone was traced manually using the polygon selection tool to define the total growth cone area. Within this region of interest, the dynamic area was segmented by applying the Huang automatic thresholding algorithm^96^ to the maximum intensity projection. The resulting binary selection was restricted to the growth cone ROI using the “Create Selection” function, and its area was measured. The dynamic area percentage was calculated by dividing the thresholded dynamic area by the total growth cone area multiplied by 100, yielding values between 0% and 100%.

Videos acquired using 2P microscopy were stabilised and deconvolved using Huygens (v25.10; Scientific Volume Imaging) prior to analysis as described above.

#### Cytoskeleton dynamics

Cytoskeleton dynamics were analysed using Fiji (v2.14.0)^95^. Maximum intensity projections were generated to quantify actin retrograde flow *in situ,* actin protrusion velocity *in vivo*, and microtubule growth speed. Kymographs were manually generated using the line tool in Fiji by drawing a line (11–25 pixels) from the tip of the growth cone to the axon shaft. Kymographs were produced using the KymoResliceWide plugin.

Actin protrusion length was measured manually in Fiji. Growth cone extensions ≥1 µm in length were classified as protrusions. To quantify microtubule fluorescence intensity from the leading edge, a line scan (11–25 pixels) was drawn from the growth cone edge to the axon shaft using the line tool. EB3 fluorescence intensity along the line was measured and normalised to the maximum intensity of each growth cone.

#### Time-integrated cross-correlation (TICCO)

Time-integrated cross-correlation (TICCO) was computed using the Pearson correlation coefficient between two image channels in a four-dimensional space (three spatial dimensions and time). Correlation values were calculated on a per-pixel basis by incorporating each pixel’s local spatial and temporal neighbourhood. The neighbourhood size was defined by the parameters *dyx* and *dt* and included all pixels along the z dimension. Accordingly, the image tile defining the neighbourhood had dimensions of (*2* × *dyx + 1*) pixels in *x* and *y*, (*2* × *dt + 1*) frames in time, and the full extent of the *z-*axis.

Prior to correlation analysis, each image tile was smoothed using a three-dimensional Gaussian filter in *x*, *y,* and *z*. The degree of smoothing in *x* and *y* was defined by *σyx,* and smoothing along z by *σz*. To avoid edge effects introduced by smoothing, the tile was increased by *2* × *σyx* in *x* and *y*. *σ* values were always chosen to be substantially smaller than *dyx* to prevent identical pixel values that would result in undefined spatial correlation. Following smoothing, tiles were cropped back to their original size for correlation calculation.

A correlation coefficient of +1.0 for a certain pixel indicates identical structures in both channels (independent of absolute intensity) that move in the same direction. A coefficient of −1.0 indicates inverse local structures (i.e., dark structures in one channel correspond to bright structures in the other) that nevertheless move in the same direction. Values close to 0 indicate the absence of correlated structures and motion. For visualisation, correlation values were inverted such that positive correlations were displayed as positive values and negative correlations as negative values.

#### CPN soma migration

CPN soma migration was analysed using Fiji (v2.16.0). Acquired images were thresholded to retain signal predominantly from cell bodies. The cortical slice was subdivided into four bins: cortical plate (CP), intermediate zone (IZ), subventricular zone (SVZ) and ventricular zone (VZ), spanning the full cortical slice. For each region, relative fluorescence intensity was quantified and expressed as a percentage of the total fluorescence intensity across the cortex.

## Statistics and figures

Statistical analyses were performed using Prism (v10.0; GraphPad) and MATLAB (vR2025a; MathWorks). Figures and schematic illustrations were assembled using Adobe Illustrator (v29.6.1; Creative Cloud).

## Acknowledgements

We thank S. Dupraz, S. A. Fiorenza, A. Husch, E. Kiemaier, A. Koseska, T. C. Lin, C. Mason, T. Misgeld, M. Oberländer, T. Pietralla, J. Schiweck, M. Sixt, S. Stern, V. Štimac, G. Tavosanis, A. Vohn, D. Wachten, R. Wedlich-Söldner, and W. Witke for reading and discussing the manuscript, M. L. Serradas for the illustrations used in the figures, C. Möhl for writing the TICCO analysis code, B. Randel, J. Gonyer, the DZNE’s Light Microscope Facility (LMF), Animal Facility (ARF), and the Collaborative Accelerator for Development and Research Engineering (CADRE) for their technical assistance.

## Author Contributions

Conceptualisation: E.A. and F.B. conceived the project and designed experiments.

Methodology: E.A., S.F. and B.E. performed the experiments.

Investigation: E.A. analysed the data.

Generation of transgenic mice: M.J.H.

Writing: E.A. and F.B.

Supervision of the project: F.B.

## Supplementary Videos

**Supplementary Video 1.**

**Axon dynamics in the intermediate zone (IZ) and corpus callosum (CC)**

Overnight tiled imaging of CPN axons in the IZ and CC *in situ*. Neurons were labelled via IUE using Lyn-mNeonGreen. P = pausing. Arrowheads mark the axon tip. Time, hh:mm; scale bars, 20 µm.

**Supplementary Video 2.**

**Growth cone morphology changes from fan-like to spike-like in the IZ-L and back to fan-like in the contralateral IZ**

Tiled time-lapse imaging of neurons labelled via IUE using Lyn-mNeonGreen. Arrowheads mark the axon tip. Dashed black lines mark the boundary between the zones. Time, hh:mm; scale bars, 50 µm.

**Supplementary Video 3.**

**Axon growth speed remains constant before, during and after turning events in the IZ-L**

A representative time-lapse from a tile scan of axons turning in the IZ-L. Neurons were labelled via IUE using Lyn-mNeonGreen. Arrowheads mark the axon tip. Time, hh:mm; scale bar, 50 µm.

**Supplementary Video 4.**

***In vivo* imaging platform (AxoVivo) used to visualise neurons in the brain of developing mouse embryos**

A multilayer setup for imaging mouse embryos *in vivo.* The intact uterine horns are placed inside the bath, which is filled with pre-warmed aCSF (37°C). Embryos are then carefully removed from the uterine sac, maintaining the umbilical cord connection, and fixed using an adjustable iris and a headplate. Finally, a coverslip is glued on top of the head. The temperature in the bath is maintained using a custom-made heating element.

**Supplementary Video 5.**

**Cortical blood flow is undisturbed for over 6 h of *in vivo* imaging in mouse embryos using AxoVivo**

Cortical blood flow was recorded using epi-fluorescence imaging. Videos were captured over 6 hours at 2-h intervals. Time, mm:ss.

**Supplementary Video 6.**

**Fan-like and spike-like growth cones have similar relative dynamic areas *in vivo***

Relative dynamics of fan-like and spike-like growth cones *in vivo* visualised using Lyn-mGreenLantern. Time, mm:ss; scale bar, 2 µm.

**Supplementary Video 7.**

**Actin distribution in the P- and C-domains is different in fan-like and spike-like growth cones *in situ***

Actin dynamics in fan-like and spike-like growth cones labelled via IUE using Lifeact-mNeonGreen (F-actin). Time, mm:ss; scale bar, 5 µm.

**Supplementary Video 8.**

**Microtubules reach further into the leading edge of spike-like growth cones *in situ***

Microtubule dynamics in growth cones labelled via IUE using Lyn-mScarlet and EB3-mNeonGreen (microtubule plus ends). Time, mm:ss; scale bar, 5 µm.

**Supplementary Video 9.**

**Polarised neurons revert back to a multipolar/neurite-deficient state following microtubule depolymerisation with nocodazole**

A representative time-lapse from a tile scan of microtubule depolymerisation in axons *in situ* using 5 µM nocodazole. Neurons were labelled via IUE using Lyn-mNeonGreen. Time, hh:mm; scale bar, 20 µm.

**Supplementary Video 10.**

**Enhanced microtubule severing by katanin overexpression inhibits axon formation *in situ***

A representative time-lapse from a tile scan of microtubule severing via katanin. Neurons were labelled via IUE using katanin-mGreenLantern. Time, hh:mm; scale bar, 10 µm.

**Supplementary Video 11.**

**Enhanced microtubule severing by spastin overexpression inhibits axon formation *in situ***

A representative time-lapse from a tile scan of microtubule severing via spastin. Neurons were labelled via IUE using spastin-mGreenLantern. Arrowhead marks the soma. Time, hh:mm; scale bar, 10 µm.

**Supplementary Video 12.**

**Enhanced microtubule stabilisation using taxol increases axon growth speed and reduces pausing *in situ***

A representative time-lapse from a tile scan of microtubule stabilisation using taxol. Neurons were labelled via IUE using Lyn-mGreenLantern. P = pausing. Arrowhead marks the axon tip. Time, hh:mm; scale bar, 20 µm.

**Supplementary Video 13.**

**Axon growth is undisturbed following blebbistatin treatment *in situ***

Tile scan of myosin II inhibition in axons *in situ* using 40 µM blebbistatin. Neurons were labelled via IUE using Lyn-mNeonGreen. Time, hh:mm; scale bar, 50 µm.

**Supplementary Video 14.**

**Movement of fibroblasts, and not growth cones, positively correlates with the movement of the ECM *in situ***

SUSHI imaging of fibroblast/growth cone interactions with the ECM. eGFP-expressing fibroblasts were transplanted into the cortex before brain sectioning. Neurons were labelled via IUE using Lyn-mNeonGreen. ECM was labelled using Alexa Fluor 647. Manual tracings of the fibroblast and growth cones from the first video frame are overlaid on the SUSHI channel and TICCO heatmaps. TICCO was used to analyse the interaction between fibroblasts/growth cones and the ECM. Scale bars, 10 µm.

---

**Extended Data Fig. 1.**
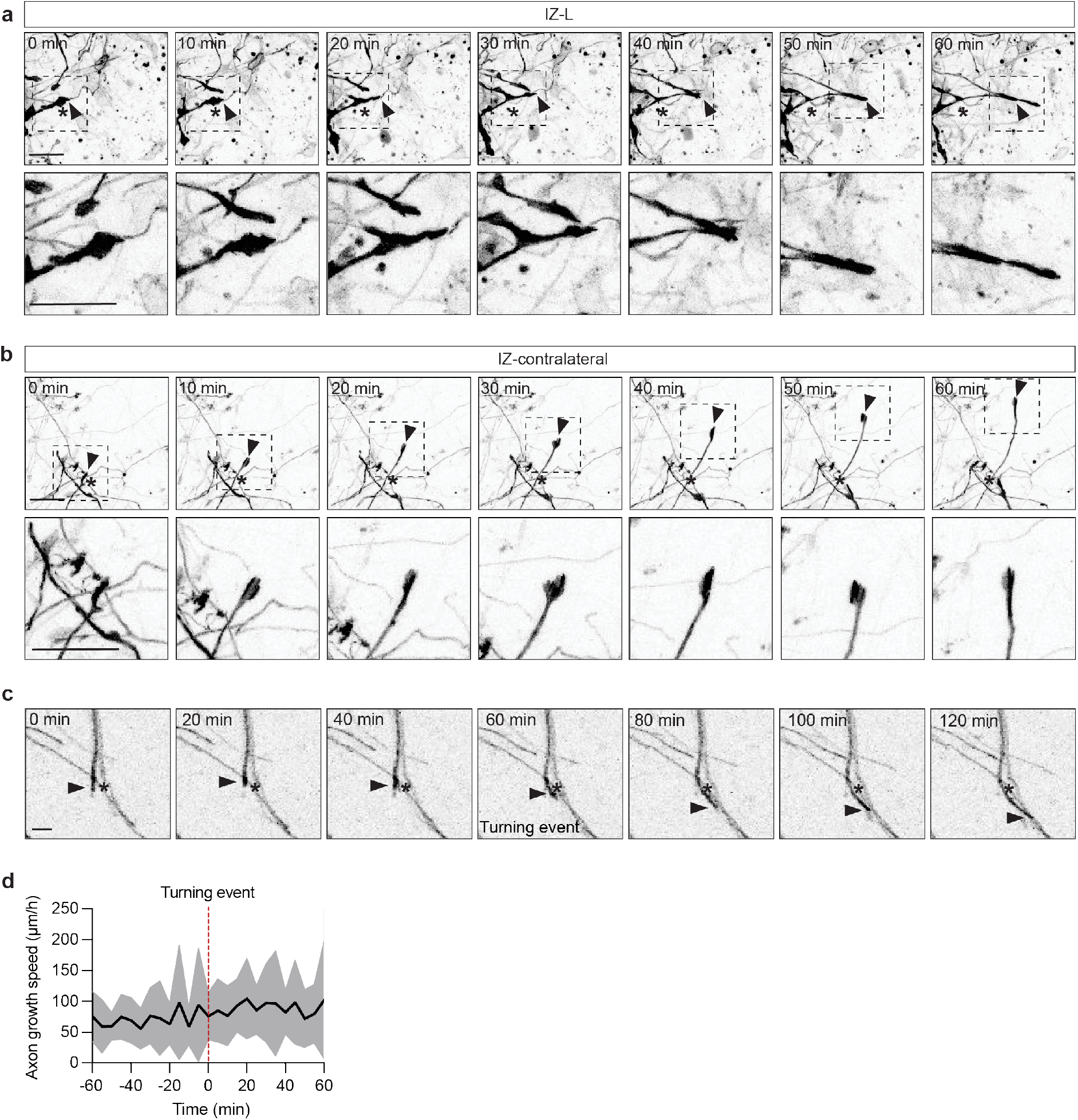
Axons undergo reversible growth cone morphological transitions and maintain growth speed during turning. a. Representative images from a tiled time-lapse of a growth cone in the IZ-L. Arrowheads mark the growth cone. Asterisks mark the initial position of the growth cone. Neurons are labelled with Lyn-mNeonGreen via IUE. Scale bars, 20 µm. b. Representative images from a tiled time-lapse of a growth cone in the contralateral IZ. Arrowheads mark the growth cone. Asterisks mark the initial position of the growth cone. Neurons are labelled with Lyn-mNeonGreen via IUE. Scale bars, 20 µm. c. Representative images from a tiled time-lapse showing an axon turning in the IZ-L. Asterisks mark the initial position of the growth cone. Arrowheads track the moving axon tip. Neurons are labelled with Lyn-mNeonGreen via IUE. Scale bar, 20 µm. d. Quantification of axon speed at turning points in the IZ-L. N = 3 independent experiments, n = 26 axons. Values are represented as a line plot with SD indicated (grey labelling).

**Extended Data Fig. 2.**
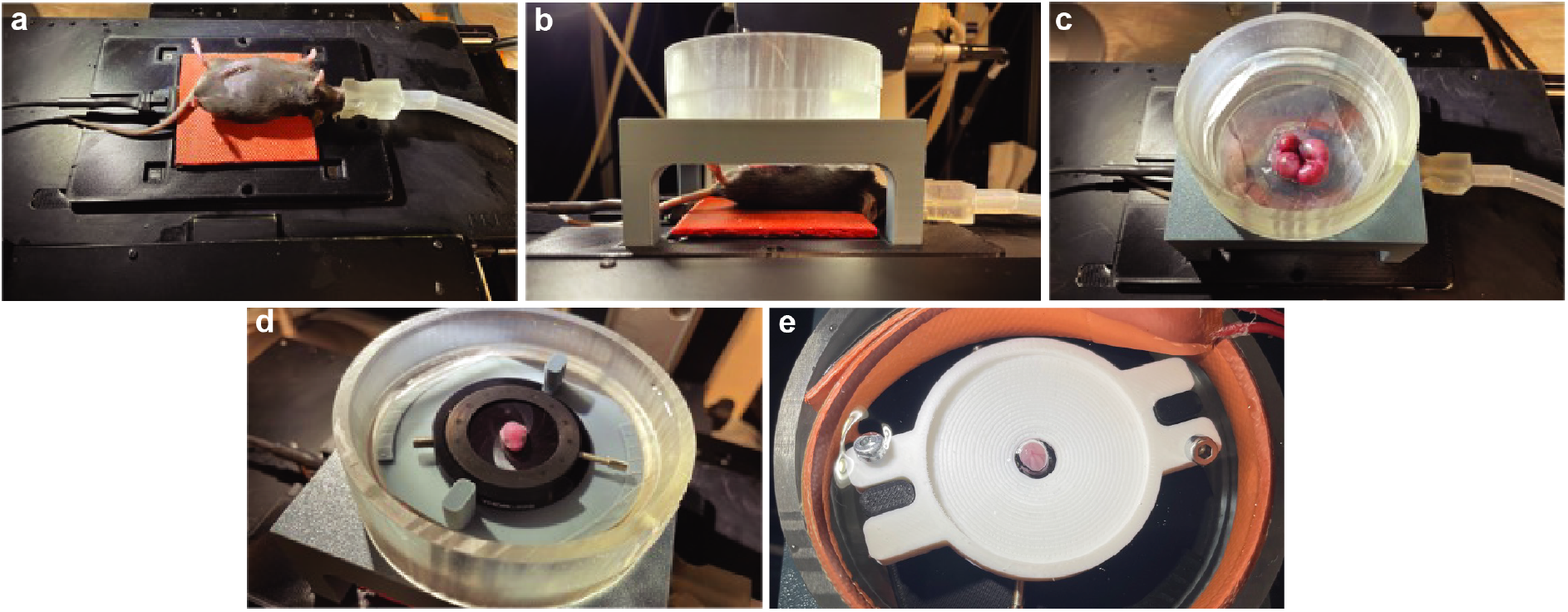
AxoVivo imaging platform enables subcellular-resolution imaging in living mouse embryos with maintained blood supply. a. An isoflurane-anaesthetised pregnant female mouse on a heating pad (37 °C). The embryos were electroporated *in utero* 2 days prior. b. A side view of the anaesthetised mouse and the imaging chamber. c. A top view of the embryos in the uterine horn placed inside the bath. d. One embryo removed from the uterus (with the umbilical cord attached) fixed with an adjustable iris. e. A top view of the embryo, immobilised using a headplate. The bath is then filled with aCSF, and the temperature is maintained at 37 °C using a heating element.

**Extended Data Fig. 3.**
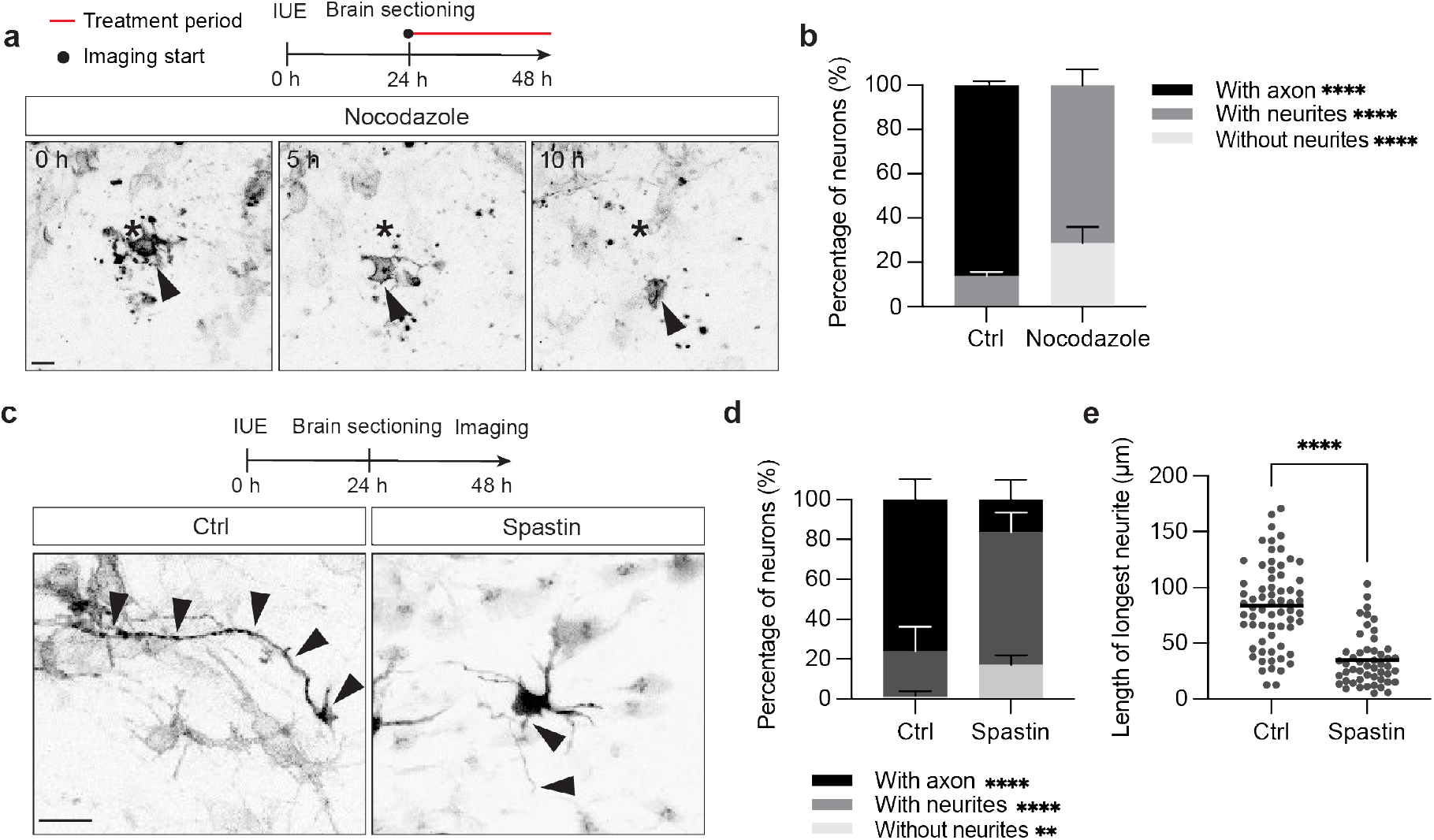
Pharmacological and molecular perturbations confirm microtubule requirement for axon formation and maintenance. a. Top, experimental design. Bottom, representative images from a tiled time-lapse showing the dynamics of a neuron treated with nocodazole *in situ*. Arrowheads indicate the soma. Asterisks mark the initial position of the soma. Neurons are labelled with Lyn-mNeonGreen via IUE. Scale bar, 10 µm. b. Percentage of neurons with an axon, with neurites or without neurites, 15 h following nocodazole treatment. N = 3 independent experiments. n = 108 neurons (control), n = 117 neurons (nocodazole). ****p <0.0001 by two-tailed Fisher’s exact test with Bonferroni correction for multiple comparisons (adjusted significance threshold α = 0.017). Each cell category (with axon, with neurite, without neurite) was compared independently between control and nocodazole using 2 × 2 contingency tables. Values are represented as stacked bars with SD indicated. c. Top, experimental design. Bottom, representative images from a tiled time-lapse of neurons expressing either spastin-mNeonGreen or only the empty plasmid containing mNeonGreen (control) via IUE. Arrowheads label the longest neurite. Scale bar, 10 µm. d. Percentage of neurons with an axon, with neurites or without neurites. N = 3 independent experiments. n = 67 neurons (control), n = 61 neurons (spastin). ****p <0.0001, **p <0.01 by two-tailed Fisher’s exact test with Bonferroni correction for multiple comparisons (adjusted significance threshold α = 0.017). Each cell category (with axon, with neurite, without neurite) was compared independently between control and spastin using 2 × 2 contingency tables. Values are represented as stacked bars with SD indicated. e. Length of the longest neurite in control and spastin-overexpressing neurons in (d). N = 3 independent experiments. n = 66 neurites (control), n = 50 neurites (spastin). ****p <0.0001 by two-tailed Mann-Whitney test. Values are represented as a scatter plot with the mean indicated.

**Extended Data Fig. 4.**
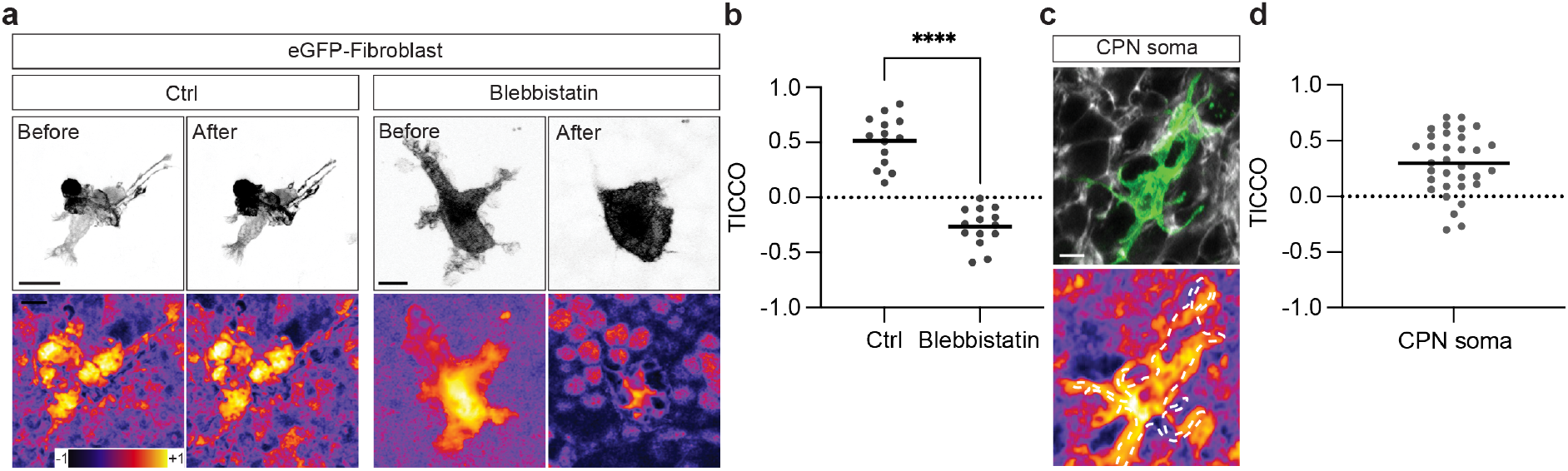
SUSHI–TICCO controls confirm sensitivity of mechanical coupling detection. a. Representative images of control and blebbistatin-treated fibroblasts (top) and TICCO-generated heatmaps (bottom). Scale bars, 10 µm. b. Quantification of ECM–fibroblast movement correlation in (a). N = 3 independent experiments, n = 14 fibroblasts (control), n = 14 fibroblasts (blebbistatin). ****p >0.0001 by two-tailed Welch’s unpaired t-test. Values are represented as a scatter plot with the mean indicated. c. A representative image of a CPN soma imaged using SUSHI (top), and a heatmap generated via TICCO analysis (bottom). Manual tracing of the soma is overlaid on the heatmap (white dotted tracing). Scale bar, 5 µm. d. Quantification of CPN soma–ECM movement correlation, analysed using TICCO. N = 4 independent experiments, n = 33 cell bodies. Values are represented as a scatter plot with the mean indicated

